# An array of signal-specific MoYpd1 isoforms determines full virulence in the pathogenic fungus *Magnaporthe oryzae*

**DOI:** 10.1101/2023.09.21.558925

**Authors:** Sri Bühring, Antonia Brunner, Klemens Heeb, Marius-Peter Mergard, Greta Schmauck, Stefan Jacob

**Affiliations:** Institute of Biotechnology and Drug Research gGmbH (IBWF), Hanns-Dieter-Hüsch-Weg 17, 55128 Mainz, Germany; Johannes Gutenberg-University Mainz, Microbiology and Biotechnology at the Institute of Molecular Physiology, Hanns-Dieter-Hüsch-Weg 17, D-55128 Mainz, Germany

**Keywords:** Ypd1, HPt protein, Alternative Splicing, Isoforms, Signal Transduction, *M. oryzae*, HOG-Pathway, ddPCR, PacBio Iso-seq, Illumina Sequencing, Virulence, Fungal Pathogen

## Abstract

*Magnaporthe oryzae* is placed first on a list of the world’s top ten plant pathogens with the highest scientific and economic importance. The pathogen is highly evolved to sense environmental changes quickly to invade different tissues of its host plant successfully. Here, we found alternative splicing (AS) as a previously unknown mechanism of how the pathogen may handle rapidly changing situations during *in planta* growth. The AS facilitates the production of multiple mRNAs and, consequently, protein isoforms with distinct biological functions from a single gene. The locus MGG_07173 occurs only once in the genome of *M. oryzae* and encodes the phosphotransfer protein MoYpd1p, which plays an important role in the high osmolarity glycerol (HOG) signaling pathway for osmoregulation. Originating from this locus, at least three *MoYPD1* isoforms are produced in a signal-specific manner. The transcript levels of these *MoYPD1-*isoforms were individually affected by external stress, which is equally present in host plant tissues. Potassium salt (KCI) stress raised *MoYPD1_T0* and *MoYPD1_T2* abundance, whereas osmotic stress by sorbitol elevates *MoYPD1_T1* levels. In line with this, signal-specific nuclear translocation of green fluorescent protein-fused MoYpd1p isoforms in response to stress was observed. The protein isoforms MoYpd1p_T0 and MoYpd1p_T2 were detected in the nucleus following KCl treatment, whereas MoYpd1p_T1 did likewise upon sorbitol stress. Mutant strains that produce only one of the MoYpd1p isoforms are less virulent, suggesting a combination thereof is required to invade the host successfully. Using *in silico* protein-protein interaction studies, we were able to assign different roles to respective isoforms in the HOG signaling pathway and predicted interactions with other signaling pathways. In summary, we demonstrate the signal-specific production of MoYpd1p isoforms that individually increase signal diversity and orchestrate virulence in the rice blast fungus *M. oryzae*.

**Authors summary:** The world’s population rises rapidly and a major problem is the global food security. *Magnaporthe oryzae* is placed first on a list of the world’s top ten plant pathogens with the highest scientific and economic importance since it causes blast, the most devastating disease of cultivated rice, which is the major food source for more than half of the world population. In this study, we demonstrate signal-specific production of a signaling protein, MoYpd1p, and show for the first time that the isoforms individually increase signal diversity and orchestrate virulence in the rice blast fungus *M. oryzae*. These results will help to better understand the molecular basis of pathogenicity of the rice blast fungus and consequently open the door to develop new strategies for plant protection and food security. In addition, our study is a valuable contribution to the basic research area of signaling mechanisms in in filamentous pathogenic fungi, and will thus be of high interest for a broad readership of the scientific community.

## 1. Introduction

*Magnaporthe oryzae* is a filamentous phytopathogenic fungus which is extremely destructive to the cultivated crop rice (*Oryza sativa*) and ranks number one among the most important plant pathogens worldwide (Dean et al. 2012). The facultative pathogen is hemibiotrophic, changing its lifestyle from bio-to necrotrophy during host invasion (Ibrahim et al. 2021). Its importance is highlighted based on the fact that almost half of the world’s population needs rice as a major food source. Even though the rice blast disease is being combated with great intensity, the pathogen annually still destructs crops that could feed more than 60 million people (Yan et al. 2023). In this respect, a better understanding of this pathogen is a prerequisite to face the global food supply. *M. oryzae* is a wonderful organism to study the molecular basis of pathogenicity, since genome and transcriptome sequences of multiple strains are available and its genome is quite suitable for directed genetic manipulation. Although the significance of alternative splicing (AS) in the rice blast fungus has already been highlighted in a few studies, details about the function of individual isoforms regarding virulence have not yet been described. All statements on virulence to date have been made based on mutants in which only the entire genomic sequence of the genes of interest have been deleted (Franceschetti et al. 2011; Li et al. 2020; Mohanan et al. 2017). There are no studies in which mutants have been created that produce only single individual isoforms.

Transcript identification and gene expression quantification have become core activities in molecular biology since the discovery of RNA’s key role as an intermediate between the genome and the proteome (Conesa et al. 2016). After the transcription of genomic DNA in eukaryotes, a protein complex called spliceosome removes introns from pre-mRNA and joins adjacent exons processing the mature mRNA (Kelemen et al. 2013; Muzafar et al. 2021; Stamm et al. 2004). Based on one genomic sequence, the spliceosome produces different mRNA transcripts through AS, which differ in stability, localization and coding sequences (CDS) (Muzafar et al. 2020; Wright, Smith, and Jiggins 2022). Consequently, several protein-coding transcript variants with different or even opposite functions can be produced from a single gene. Therefore, AS is a fundamental mechanism in eukaryotes and enhances the regulatory and functional diversity of proteins and phenotypic traits (Bush et al. 2017; Wright et al. 2022). The diversity of AS events are classified into five main categories (Figure 1): the removal of a single exon (exon skipping), the retention of one intron in the mRNA (intron retention), alternative 5′ or 3′ splice sites and mutually exclusive exons.

**Figure 1:**
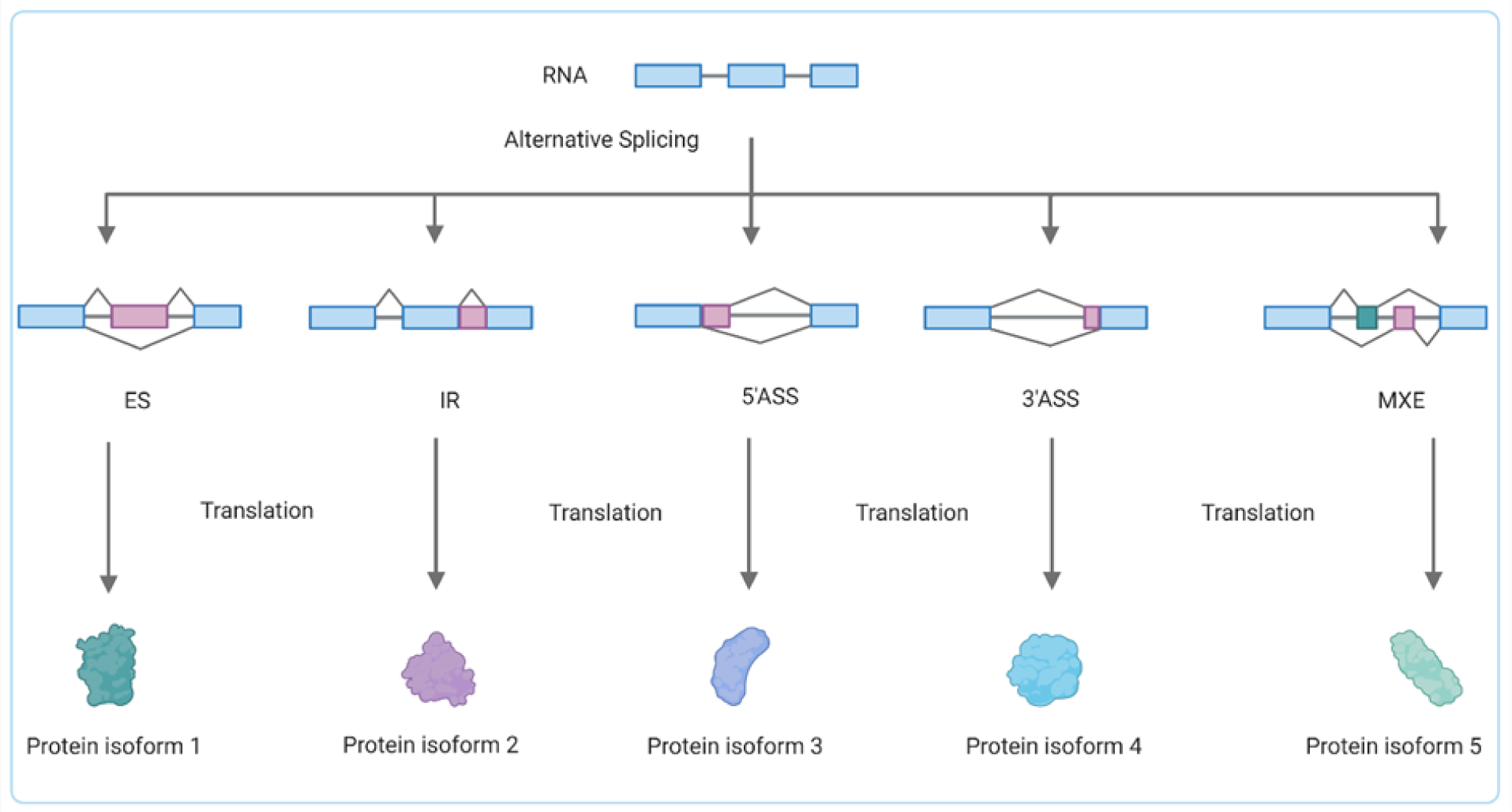
Simplified illustration of the five major alternative splicing (AS) patterns. The exons are represented as boxes and the introns as lines. The AS of the pre-mRNA via exon skipping (ES), intron retention (IR), alternative 5′ or 3′ splice sites (5′ ASS, 3′ ASS) and mutually exclusive exons (MXE) can produce different proteins.

Intron retention is the most commonly described AS event in fungi (Fang et al. 2020; Grützmann et al. 2014; Jeon et al. 2022). Even though the AS phenomenon has been known for over 40 years, the annotated transcripts only partially reflect the huge complexity of all AS events in fungi. Missing or incorrect genome annotations and the availability of datasets only about the most popular gene transcripts in the fungal databases limit our current knowledge of the biological impact of AS (Sieber et al. 2018). Increasing the analysis of RNA-seq data suggests a critical role for AS in the fungal kingdom (Fang et al. 2020; Grützmann et al. 2014; Jeon et al. 2022). The histidine-containing phosphotransfer protein Ypd1p plays an important role in osmoregulation as part of the high osmolarity glycerol (HOG) pathway. The latter is also involved in the regulation of growth, development and virulence in the rice blast fungus *M. oryzae*, and overstimulated by the phenylpyrrole fungicide fludioxonil (Bersching and Jacob 2021a; Jacob et al. 2014; Zhang et al. 2021).The signaling pathway is composed of a multistep phosphorelay (MSP) system followed by a mitogen-activated protein kinase (MAPK) cascade downstream. The MSP system comprises the hybrid sensor histidine kinases (HKs) MoHik1p and MoSln1p, the histidine-containing phosphotransfer (HPt) protein MoYpd1p and the response regulator (RR) MoSsk1p. Signal phosphoryl group transfer is mediated in these MSPs by the His-Asp-His-Asp manner (Singh et al. 2021). In response to changing environmental osmolarity, MoYpd1p acts as the central linchpin and transmits signals through reversible phosphorylation and dephosphorylation within the MSP. Under isosmotic conditions, the MSP system is constitutively phosphorylated, thereby, inhibiting the MAPK cascade. By contrast, the signaling components are dephosphorylated as external osmolarity rises, which results in the activation of the MAPK cascade by phosphorylation (Jacob et al. 2015). Finally, the phosphorylated MoHog1p translocates into the nucleus and induces the stress response (Bohnert et al. 2019). Compared to most fungal genes, only one MoYPD1 gene is annotated in the genome with two transcripts (MoYPD1_T0 and MoYPD1_T1). It is well-known that MoYpd1p interacts with MoSln1p and MoHik1p in the MSP (Jacob and Thines 2017). Apart from MoSln1p and MoHik1p, eight further HKs are encoded in the genome of *M. oryzae*, which may also can perform protein-protein interactions (PPI) with MoYpd1p (Jacob et al. 2014). Moreover, a third transcript isoform *(MoYPD1_T3)* was identified on the cDNA and protein level (Bühring et al. 2021). In addition to its key role in the HOG signaling pathway, *MoYPD1* appears to positively and negatively influence genes involved in other stress adaptation processes and virulence (Mohanan et al. 2022). Studies on Ypd1p homologs in other fungi, such as *Candida albicans* and *Aspergillus fumigatus*, have shown that green fluorescent protein (GFP)-fused Ypd1p shuttles between the cytoplasm and the nucleus (Mavrianos et al. 2014; Schruefer et al. 2021). Therefore, it is conceivable that MoYpd1p interacts with targets in both the cytosol and nucleus. However, the specific function of *MoYPD1* isoforms and their potential for signal-specific functions still remains enigmatic. A better understanding of the complex role of *MoYPD1* in *M. oryzae* can be obtained by the identification and characterization of additional isoforms. We used various bioinformatics tools and validated the predictions in molecular biology experiments to answer the questions of which *MoYPD1* transcript isoforms exist and whether they are produced or function in a signal-specific manner in the HOG pathway.

## 2. Results

### 2.1 Prediction of novel interactions between MoYpd1p and histidine kinases

Our studies reveal new, previously unknown interactions between MoYpd1p and different HKs that may facilitate and explain the diversity in signal transduction in *M. oryzae* and explain its virulence. The PPI analyses based on the three-dimensional (3D) structures of the isoforms and the HKs were modulated to gain insight into the physiological functions of MoYpd1p_T0 (MGG_07173T0), MoYpd1p_T1(MGG_07173T1) and MoYpd1p_T2 (Bühring et al. 2021). Due to the unpredictable 3D model caused by the size of MoHik6p, it was impossible to calculate its interaction with isoforms. As shown in Table S3, the isoforms of MoYpd1p are possible interaction partners for all the HKs except MoHik7p. Regarding the HOG pathway, PPIs most likely occurred between MoSln1p and MoYpd1p_T1 and between MoHik1p and MoYpd1p_T0 (Figure 2, Figure S1).

**Figure 2:**
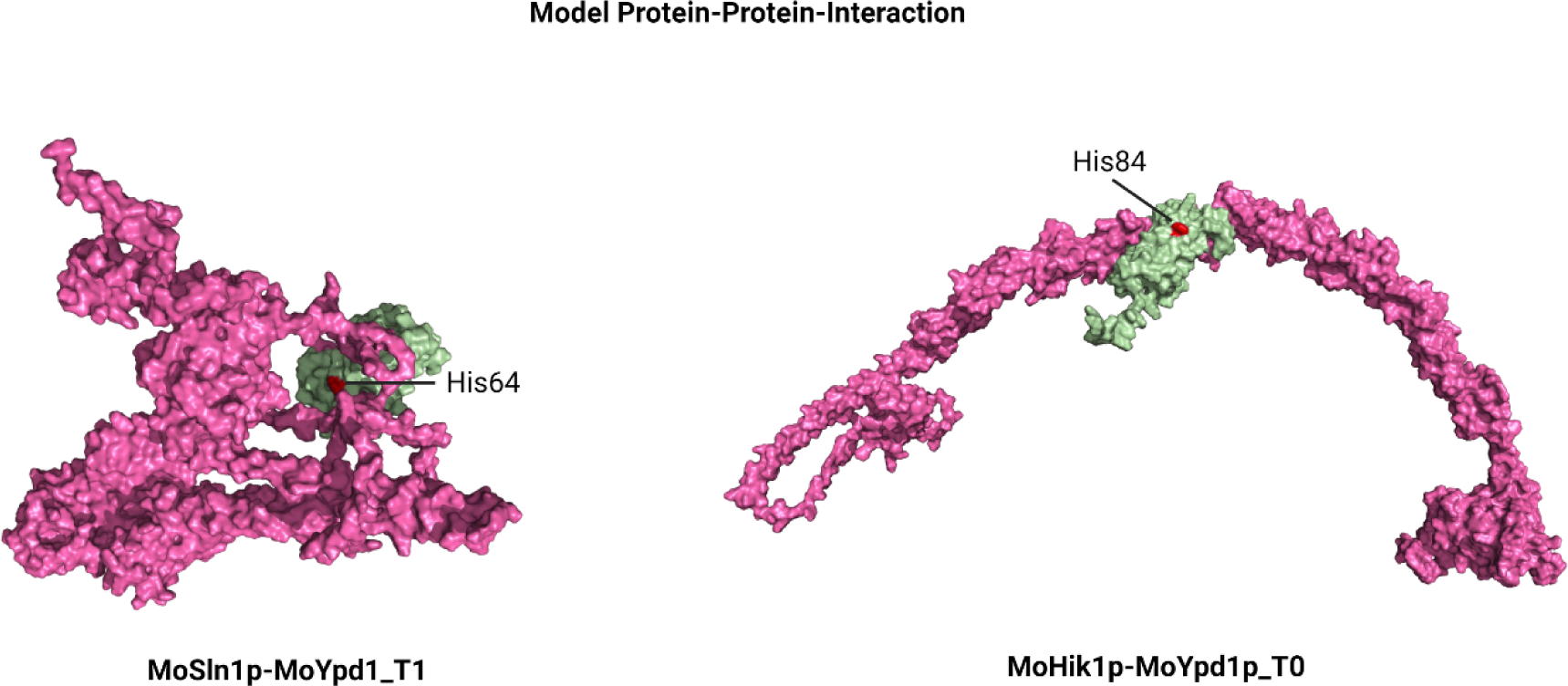
Protein-Protein Interaction. Prediction between MoYpd1p_T1 (green) and MoSln1p (pink) (A) and between MoYpd1p_T0 (green) and MoHik1p (pink) (B). Red dot: H82 (A) and H64 (B) of the MoYpd1p HPt domain involved in the interaction.

Furthermore, less pronounced interactions between MoHik1p and MoYpd1_T1 or MoSln1p and MoYpd1_T0 were calculated (Table S1). Such PPIs were not predicted for MoYpd1p_T2 (Figure S1). However, interactions of all isoforms were modulated between MoHik3p, MoHik4p and MoHik8, respectively (Figure S2). In addition, MoHik3p was found directly interacting with the isoform MoYpd1p_T2. MoYpd1p’s HPt domain contains amino acids His64 (in MoYpd1p_T0), His84 (in MoYpd1p_T1) and His92 (in MoYpd1p_T2), which have been identified as important for phosphorylation/dephosphorylation reactions during the predicted interactions between HKs and the isoforms MoYpd1p_T0–T2 tested.

### 2.2 Signal-specific localization of MoYpd1p isoforms and their role in pathogenicity

In order to identify and follow spatial localization of the different phosphotransferase isoforms, we generated different mutant strains by fusing a GFP to the genomic sequence of *MoYPD1* (referred to on a protein level as total MoYpd1p) and to isoform-specific cDNA sequences (MoYpd1p_T0, MoYpd1p_T1, MoYpd1p_T2, Supplementary, Table 4) and used the loss-of-function mutant *ΔMoypd1* as a parent strain. Consequently, we have the mutant strain *ΔMoypd1::MoYPD1-GFP* producing “all” isoforms, and the mutant strains *ΔMoypd1::MoYPD1_T0-GFP, ΔMoypd1::MoYPD1_T1-GFP* and *ΔMoypd1::MoYPD1_T2-GFP* producing only the single isoforms MoYpd1p_T0, MoYpd1p_T1 and MoYpd1p_T2, respectively.

The GFP signal in all mutant strains was observed to be distributed within the cytoplasm under isoosmotic conditions, whereas the respective subcellular localization of total MoYpd1p and the three isoforms was found in the nucleus in response to different stress stimuli. In detail, we found an accumulation of the GFP-signal of total MoYpd1p in the nuclei of the mutant strain *ΔMoypd1::MoYPD1-GFP* one minute after stress exposure to KCl [1 M], NaCl [0.75 M] or sorbitol [1 M] (Figure 3A). By contrast, the spatial localization of MoYpd1p_T0, MoYpd1p_T1 and MoYpd1p_T2 changes individually after stress, providing a deeper understanding of the role of AS in signal diversity within phosphorelay systems. The respective isoforms MoYpd1p_T0 and MoYpd1p_T2 translocate into the nucleus in the mutant strains *ΔMoypd1::MoYPD1_T0-GFP* and *ΔMoypd1::MoYPD1_T2-GFP* as a result of 0.75 M KCl stress, whereas the isoform MoYpd1p_T1 in the mutant strain *ΔMoypd1::MoYPD1_T1-GFP* was located within the nucleus in response to sorbitol treatment (Figure 3B). After exposure to sorbitol stress [1 M], the GFP-signal of both isoforms MoYpd1p_T0 and MoYpd1p_T2 remained in the cytoplasm.

**Figure 3:**
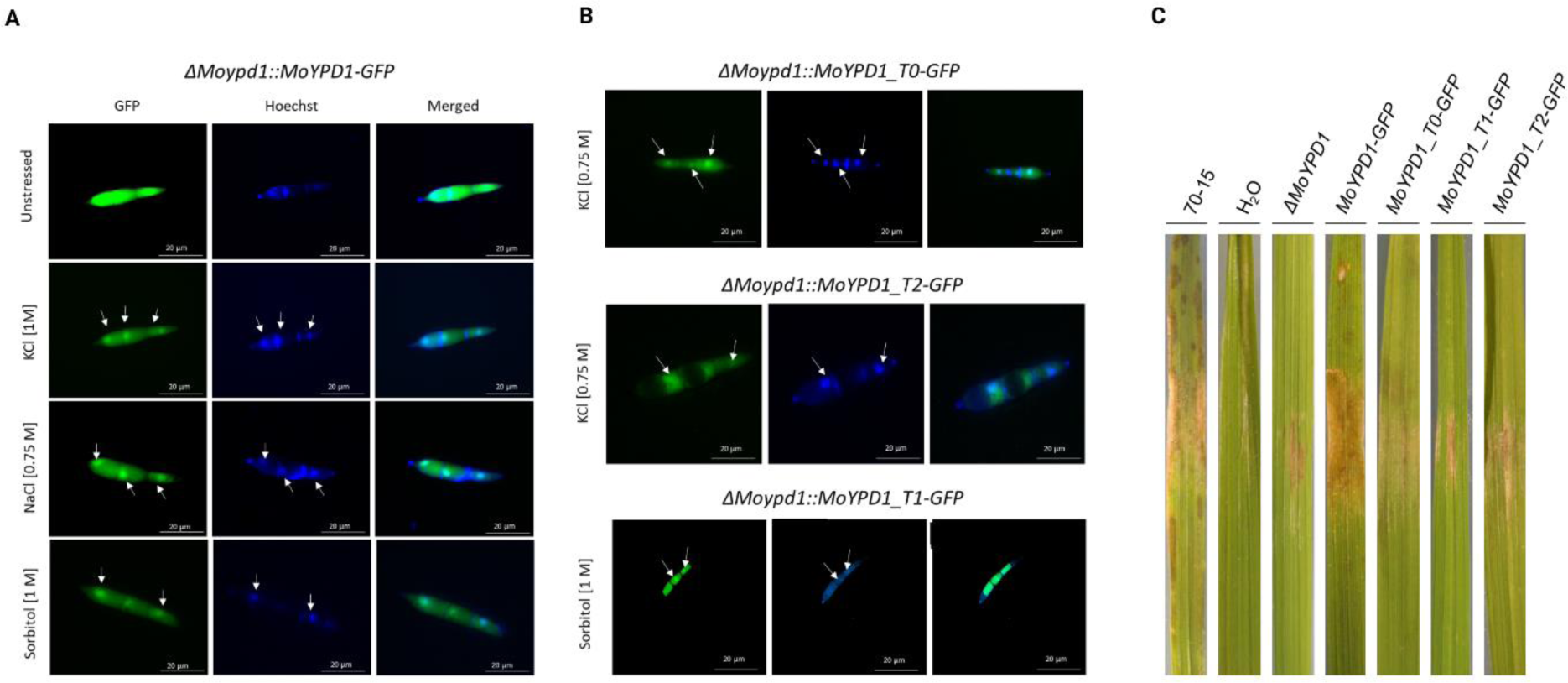
Localization of GFP-fused total MoYpd1p and different MoYpd1p isoforms. A: Translocation of GFP-fused total MoYpd1p 1 min after salt (KCl [1 M] and NaCl [0.75 M]) or sorbitol [1 M] treatment. The GFP signal in the untreated control was distributed throughout the cytoplasm of the mutant strain *ΔMoypd1::GFP-MoYPD1*. B: Localization of GFP-fused MoYpd1p isoforms. After treatment with KCl [0.75 M], the translocation into the nucleus was observed for MoYpd1p_T0 and MoYpd1p_T2. However, MoYpd1p_T1 was found to accumulate in the nucleus upon sorbitol stress [1 M].C: Lesions on rice leaf surfaces three days after inoculation with *M. oryzae*. Detached rice leaves were inoculated with a conidial suspension of the wildtype strain 70-15, *ΔMoypd1::MoYPD1-GFP*, *ΔMoypd1::MoYPD1_T0-GFP*, *ΔMoypd1::MoYPD1_T1-GFP* or *ΔMoypd1::MoYPD1_T2-GFP*. Mutant strains producing only one of the isoforms were found to cause fewer disease symptoms on the leaves compared to the control strains.

The role of MoYpd1p and its isoforms in virulence is shown in Figure 3C. The wildtype strain 70-15 was found to cause significant lesions on detached rice leaves, whereas the mutant strains producing only one of the isoforms MoYpd1p_T0, MoYpd1p_T1 or MoYpd1p_T2 have been documented to be less virulent. The positive control of the complemented mutant strain *ΔMoypd1*::*MoYPD1-GFP* shows almost as strong infection symptoms as those in the wildtype strain, indicating that all isoforms collectively are needed for full virulence in *M. oryzae*. Experiments on whole rice plants inoculated with conidial suspensions underlined this hypothesis. Similar to the detached leaf assays, the same results could be observed in the plant assays (Figure S4A).

In line with this, a vegetative growth assay validated the need for an interplay of all MoYpd1 isoforms in order to achieve a completely functional osmoregulation in *M. oryzae* (Figure S4B). MoYpd1p-producing isoform strains showed growth inhibition of over 70 % in the presence of high concentrations [0.4 and 0.6 M] of KCl or sorbitol, suggesting that the role of the individual isoforms is insufficient to cope with conditions of sustained high osmolarity.

### 2.3 Additional *MoYPD1* isoforms and signal specific splicing

As a result of our replicate multivariate analysis of transcript splicing (rMATS) analysis, we were able to determine the total number of AS events of protein-coding genes in *M. oryzae* between the unstressed condition and KCl [0.5 M], sorbitol [0.5 M] and fludioxonil-stressed [10 µg/ml] conditions. In this context, 3’ ASS (about 37 %) and 5’ ASS (approximately 30 %) were identified as the most common AS patterns in all samples. The total number of AS events after exposure to KCl increased from 3474 to 3851. The number of AS events over the same period after sorbitol exposure were found to be decreased from 3661 to 3292, and from 3474 to 2766 after treatment with fludioxonil (Figure 4A, Table S5). How the stress conditions tested influence MoYPD*1* splicing patterns is shown in Sashimi plots (Figure 4B).

**Figure 4:**
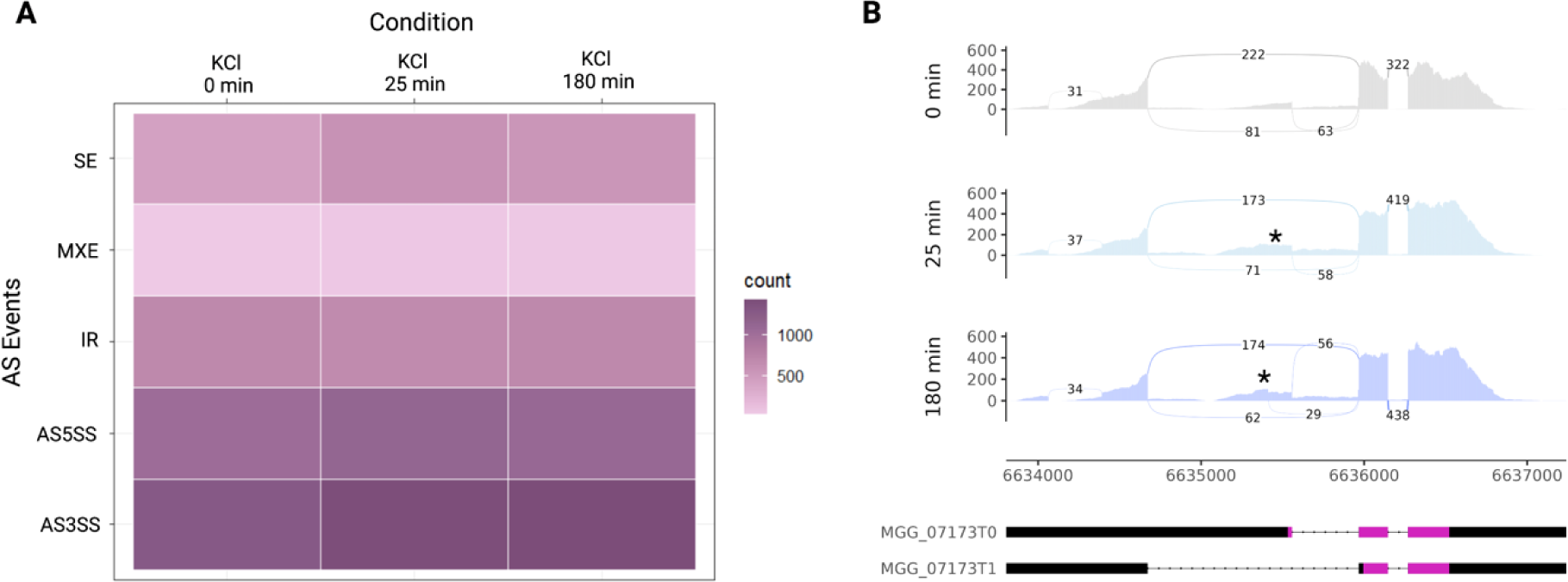
Evaluation of RNA-seq data. A: Heat map of RNA-Seq transcriptome analysis for AS Events from *M. oryzae* under unstressed and stressed conditions (25 and 180 min after treatment with 0.5 M KCl. 3’ ASS and 5’ ASS were found to be the most common AS patterns following KCl stress induction. B: Sashimi plot visualized the AS pattern of *MoYPD1* during 0.5 M KCl stressed and unstressed conditions. In addition to the annotated introns of MGG_07173T0 and MGG_07173T1, additional splice junctions are shown (see asterisk).

Coverage tracks are illustrated in gray and purple, respectively, with exons connected by arcs indicating splicing events. Annotated transcripts of (isoform MoYpd1p_T0 and M (isoform MoYpd1p_T1 are presented below the plots, where pink represents the CDS. In addition to the stress-independent splicing events of the annotated isoforms and an additional splice junction (Chr2:6634064-6634391) in the 5’ direction of the CDS, unannotated splice sites were also revealed by KCl exposure (marked by an asterisk). A similar splicing pattern was observed upon sorbitol and fludioxonil-induced stress (Figure S4 and S5). Based on our rMATS analysis, four possible new *MoYPD1* isoforms with alternative splice regions have been identified. We detected the isoforms using event- and isoform-based prediction tools and underpinned this with an IsoSeq-based transcript assembly (Table 1).

**Table 1:** Selection of different *MoYPD1* transcript isoforms using different prediction tools.

| Tools | Isoform T0<br>Chr2:6633806–6635558 | Isoform T1<br>Chr2:6633806–6634673 | Isoform T2<br>Chr2:6633806–6635020 | Isoform T3<br>Chr2:6633806–6635879 | Isoform T4<br>Chr2:6633806–6634673 | Isoform T5<br>Chr2:6633806–6635411 |
| --- | --- | --- | --- | --- | --- | --- |
| rMATS | ✓ | ✓ | ✓ | ✓ | ✓ | ✓ |
| Cufflinks | ✓ | ✓ | – | – | ✓ | – |
| IsoSeq | ✓ | ✓ | – | – | ✓ | – |
| MAJIQ+Voila | ✓ | ✓ | – | ✓ | ✓ | ✓ |
| SGSeq | ✓ | ✓ | – | ✓ | ✓ | ✓ |

The *in silico* analysis contributed 5’GT-AG 3’ as a splice site; this motif is known to be a critical factor for properly splicing RNA in eukaryotes. The region between the two posterior exons is a conserved region of the isoforms; differences in splice patterns result from the length of the first intron. An exception is the isoform (isoform T4) mentioned previously with an additional exon. The MoYpd1p isoforms T0, T1 and T4 were consistently detected by all methods, indicating their widespread presence and biological importance. However, even though rMATS, modeling alternative junction inclusion quantification (MAJIQ)+Voila and SGSeq predicted the existence of the transcript MoYpd1p isoforms T3 and T5, only rMATS detected the presence of MoYpd1p_T2, an isoform already determined at the peptide level (Bühring et al. 2021). After the prediction of various isoforms for *MoYPD1*, the MoYpd1p isoforms T1–T5 were amplified at the cDNA level and confirmed by sanger sequencing. The ORF prediction using NCBI ORF Finder reveals a novel ORF for isoform T3 in addition to the ORFs already known (Table S6).

Isoform expression analyses were performed with DESeq2 during the next step in order to investigate potential signal-specific functions of the isoforms. The most prevalently expressed isoform under non-stressed conditions was T1, followed by T0 and T4. According to our findings, the transcript MoYpd1p isoforms T2, T3 and T5 exhibited a consistently low-frequency expression, whereas the FPKM values of the remaining isoforms change depending on the stress induction. The expression of MoYpd1p_T0 increases after treatment with 0.5 M KCl, whereas the expression of MoYpd1p_T1 increases after treatment with 0.5 M sorbitol and 10 µg/ml fludioxonil (Figure 5A Figure S7).

**Figure 5:**
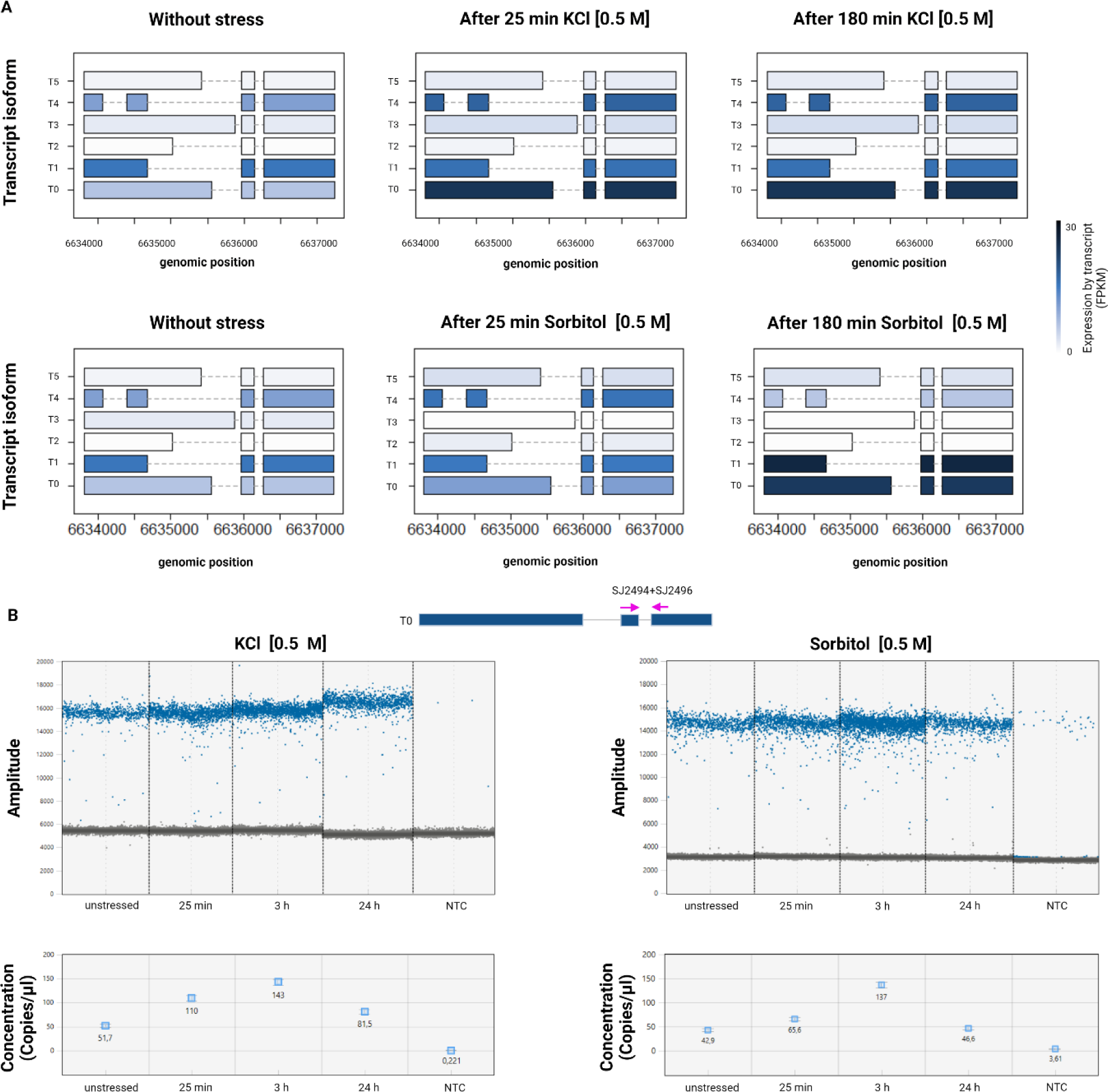
*MoYPD1* isoform expression analysis. A (Top): Heat plot of *MoYPD1* isoform expression under different conditions. Isoosmotic conditions result in the highest FPKM values for MoYpd1p_T1. The expression level of isoform T0 after 25 and 180 min of 0.5 M KCl treatment exceeds that of MoYpd1p_T1, resulting in MoYpd1p_T0 being the dominantly expressed isoform. However, the transcript level of isoform T1 is highest after sorbitol stress (Buttom). B: ddPCR result of the conserved *MoYPD1* region. Top:1 D plot of ddPCR reaction under KCl- and sorbitol-stressed conditions. A blue dot indicates a positive droplet containing at least one copy of *MoYPD1* and a gray dot indicates a negative droplet lacking any target DNA. Bottom: Concentrations (copies/μl) of the conserved *MoYPD1* region, as processed by QuantaSoftware™. The error bars show the maximum and minimum Poisson distributions for the 95 % confidence interval.

Fludioxonil strongly increases MoYpd1p_T4expression as well. In conclusion, these observations demonstrate that the isoforms T0 and T1 are dominantly produced under isoosmotic conditions. Stress induces the translocation of these proteins into the nucleus and, simultaneously, increases their expression level. Furthermore, the results are underpinned using droplet digital polymerase chain reaction (ddPCR) analysis investigating the absolute expression pattern (Figure 5B). In order to amplify all possible isoforms, we used the primer combination SJ2494 and SJ2496 (Table S7), capable of amplifying the conserved region in the gene. As a result of KCl and sorbitol stress, we observed a nearly 2- and 3-fold increase, respectively, in copies/µl at 25 and 180 min for the conserved *MoYPD1* region, which decreased pre-stress levels after 24 h. Taken together, our results confirm our hypothesis of the existence of more isoforms of *MoYPD1* than previously described. Additionally, we provide evidence for signal-specific functions of the isoforms in the transmission of KCl, sorbitol and fungicide stress, respectively.

### 2.4 A new model of the HOG pathway comprises more elements

Based on the results presented in this study, we concluded that the *MoYPD1_T0* isoform is dominantly expressed after KCl stress. By contrast, the *MoYPD1_T1* isoform was found to be expressed mainly after sorbitol stress. In accordance with our observations of nuclear translocations of the GFP-fused protein isoforms MoYpd1p_T0 and MoYpd1p_T1 upon KCl and sorbitol stress induction, respectively, we present an extension of the current model of the HOG pathway including these isoforms (Figure 6). The phosphorelay system is active in isoosmotic (untreated) conditions, thereby inhibiting the downstream MAPK cascade. Phosphorylated MoYpd1p isoforms are in the cytoplasm. Following KCl and sorbitol stress perception by MoSln1p and MoHik1p, respectively, MoYpd1p_T0 and MoYpd1p_T1 translocate into the nucleus immediately after stress induction. High osmolarity inactivates the phosphorelay system, activates the MAPK cascade and leads to phosphorylation of MoHog1p, which is thereby translocated to the nucleus. According to our findings, MoHog1p and the isoforms MoYpd1p_T0/T1 or a combination of them are probably responsible for initiating the nuclear reaction. In addition, our PPI models suggest interactions between the two isoforms MoYpd1p_T0 and MoYpd1p_T1 and MoHik3p, MoHik4p and MoHik8p. Only interactions between MoHik4p and MoHik8p were predicted for MoYpd1_T2.

**Figure 6:**
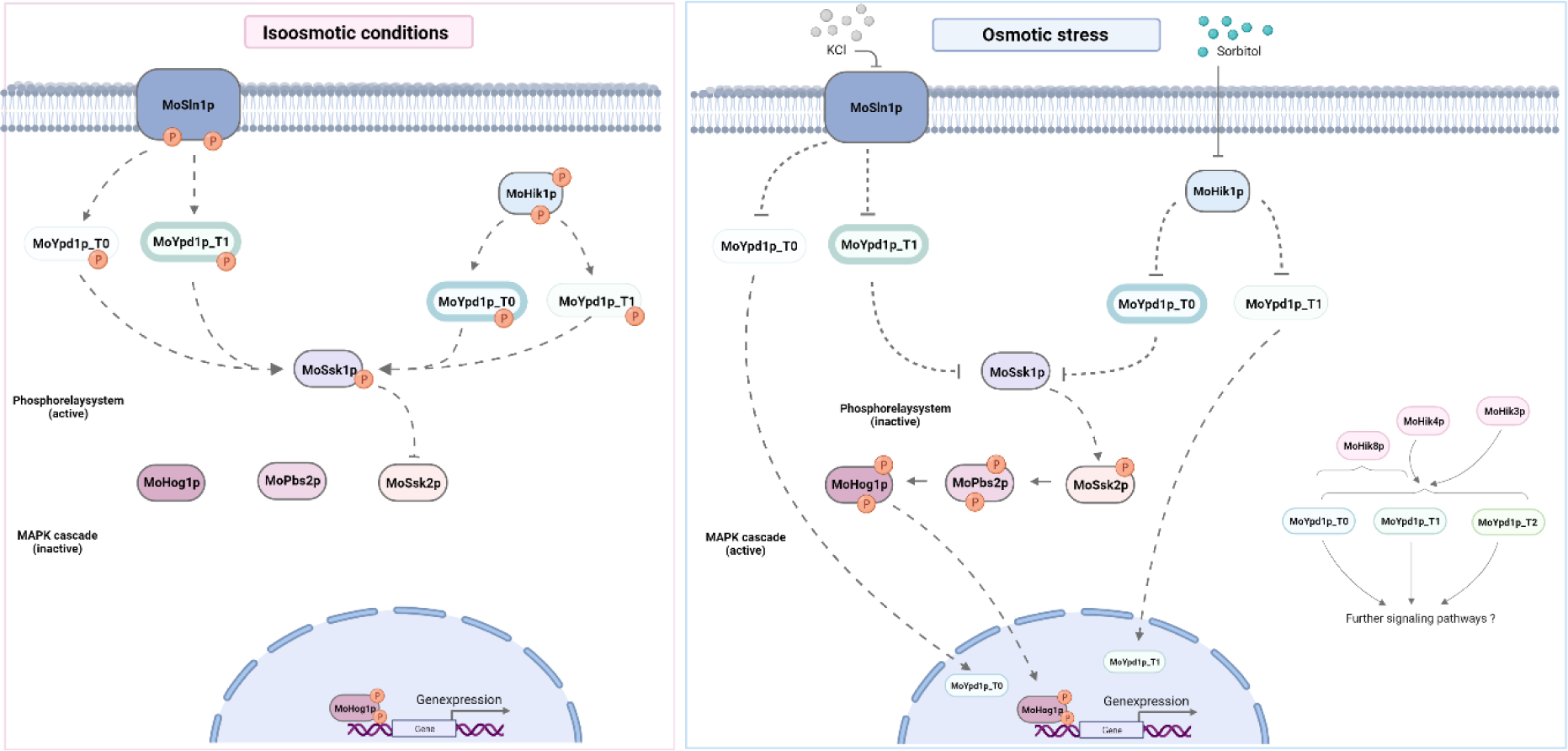
High osmolarity glycerol pathway in *M. oryzae*. MoYpd1p_T0 and MoYpd1p_T1 translocate to the nucleus after sensing KCl and sorbitol stress by MoSln1p and MoHik1p, respectively. Depending on the stress, phosphorylated Mohog1p and dephosphorylated Moypd1p_T0 or MoYpd1p_1 initiate a response in the nucleus. The figure illustrates the possible PPI between isoforms and the HK MoHik3p, MoHIk4p and MoHik8p.

## 3. Discussion

Globally, *M. oryzae* is one of the most devastating agricultural pathogens (Fernandez and Orth 2018; Nalley et al. 2016). That is down to the fact that rice blast disease is a major threat to the food security of approximately half of the world’s population who depend on rice as their primary food. A better understanding of the pathogen’s biology is essential to address this problem. Pathogenicity-related processes are shaped by differentiation, multicellular communication mechanisms, modulation or rewiring of cellular pathways, and/or suppression of defense mechanisms (Cai et al. 2021; Lai et al. 2022; Ying, Feng, and Keyhani 2013). Consequently, the development of effective control strategies based on a deeper understanding of signaling mechanisms involved in fungal virulence is a promising path in plant protection (González-Rubio et al. 2019; Jacob, Bühring, and Bersching 2022). *M. oryzae* needs to adapt rapidly to changing osmotic conditions in the plant tissue during the intracellular invasion of the host plant. Osmoregulation in fungi is coordinated via the HOG pathway (Hohmann, Krantz, and Nordlander 2007). Although the latter has been well studied in *Saccharomyces cerevisiae*, its regulatory network and contribution to the virulence of fungal pathogens remains to be elucidated (Huang et al. 2021; Yaakoub et al. 2021). The HOG pathway is also known as the target of the fungicide fludioxonil, however, its mode of action is not completely understood (Bersching and Jacob 2021b; Brandhorst et al. 2019). Based on Y2H experiments, it was found that the phosphotransferase (HPt) MoYpd1p acts as the key mediator of the phosphorelay system of the HOG pathway, orchestrating the phosphorylations between the two sensor HKs MoHik1p and MoSln1p as well as the RR MoSsk1p (Jacob et al. 2015). Signaling systems, such as phosphorelay systems, generally use proteins containing HPt to transmit signals from sensor HKs to RRs (Alvarez and Georgellis 2022; Jacob et al. 2022). Based on RNA-Seq and HPLC-MS/MS analyses, we previously hypothesized that AS may produce several different MoYpd1p isoforms with distinct physiological functions (Bühring et al. 2021; Jacob and Thines 2017). Apart from MoSln1p and MoHik1p, MoYpd1p is likely to interact with one or more of the remaining eight HKs (MoHik2p-MoHik9p) identified in *M. oryzae* (Jacob et al. 2014). Here, we support this assumption with our PPI models, which revealed new, previously unknown interactions between MoYpd1p and three HKs, thus, explaining AS as a molecular mechanism for increasing diversity in signal transduction in *M. oryzae* (Figure 2, Figure S1, Table S1). Furthermore, the aim of the present investigation was to identify novel *MoYPD1* isoforms *in silico* and determine their role in pathogen signal-processing during infection *in vivo.* Up to now, the scientific understanding of AS and the role of different isoforms during fungal infection processes and development has still been limited (Jeon et al. 2022; Muzafar et al. 2021). However, numerous studies in plants have shown that AS is a crucial posttranscriptional mechanism for regulating gene expression and produces various isoforms, thereby, controlling biotic and abiotic stress responses (Chen et al. 2019; Laloum, Martín, and Duque 2018; Mandadi et al. 2023). The RGA5 resistance gene in rice (*Oryza sativa*), for example, is transcribed into the two isoforms RGA5-A and RGA5-B, conferring resistance of rice to *M. oryzae* effector proteins AVR-Pia and AVR1-CO39 (Cesari et al. 2013; Ganie and Reddy 2021; Younas et al. 2023). Previous studies in plants have also demonstrated that distinct Hpt proteins interact differently with RR, thereby, influencing signal transduction in the HOG and cytokinin signaling pathways (Bertheau et al. 2015; Héricourt et al. 2019). Contrary to this, the effect of isoforms as virulence factors in the fungal kingdom is rare. In order to change and tackle this, we identified different isoforms of *MoYPD1* and elaborated their contribution to fungal physiology and pathogenicity in *M. oryzae*.

It is generally well accepted that signaling proteins containing Hpt, such as Ypd1p, are central constituents of pathogenicity-related differentiation and fungal virulence (Day et al. 2017; Lee et al. 2011). The first hint of the existence and involvement of multiple MoYpd1p isoforms in *M. oryzae* pathogenicity resulted from our comparison of disease symptoms on wounded rice leaves inoculated by different *Magnaporthe* mutant strains. In this study, plant assays with fungal mutants carrying single aberrant MoYpd1p isoforms highlighted, for the first time, the important role of each individual isoform acting together in a holistic machinery to obtain full virulence. Indeed, mutant strains producing just one sole MoYpd1p-isoform caused only a limited number of lesions on the rice leaves compared to the wildtype strain and the complemented mutant strain *ΔMoYpd1::MoYPD1-GFP*. These results are consistent with those of a transcriptome analysis of the *ΔMoYPD1* mutant of strain B157.Several genes involved in cell wall degradation and secondary metabolite production were found to be downregulated in this analysis (Mohanan et al. 2022).

We present here the first report about the contribution of single isoforms of a signaling protein to pathogenicity-related processes *in vivo* in fungal pathogens. Moreover, these results are consistent with limited previous studies shedding light in the general linkage of AS and fungal virulence. Disease symptoms, for example, were detected on cabbage after inoculation of the wildtype strain of *Fusarium oxysporum f. sp. conglutinans (Foc*) and a mutant strain producing four isoforms of the effector protein SIX1. However, no disease symptoms were observed in the cabbage after inoculation of the *Foc-ΔSIX1* mutant strain (Li et al. 2016). In addition, infection studies on *Candia albicans* mutants that produce protein mannosyltransferase isoforms have shown that the virulence is dependent on not only the individual isoforms but also the environment in which they are infected (Rouabhia et al. 2005). The expression profiles of the yeast *Hortaea werneckii* highlight a specific role of the MAP2K HwPbs2 isoforms within the HOG pathway. Consequently, two HwPbs2 isoforms (HwPbs2A and HwPbs2B2) are involved in the rapid adaptation to a hypersaline environment, while a third isoform (HwPbs2B1) appears to respond to moderate salt stress (Plemenitaš 2021).

Identifying isoforms and elucidating their physiological functions require synergistic approaches of molecular biology and bioinformatics. Thereby, RNA-Seq data analysis as the only basis can be problematic for several reasons. Most published fungal RNA-Seq data are not strand-specific, and transcripts frequently overlap between adjacent genes due to their high gene density (Cao et al. 2022). Furthermore, choosing between sequencing technologies and a wide range of bioinformatic tools that perform similar tasks but have different advantages and disadvantages can be challenging (Mehmood et al. 2020; Thind et al. 2021). We initially identified various *MoYPD1* isoforms using both short-read Illumina RNA-Seq and long-read PacBio Iso-seq sequencing methods, and subsequently verified their existence at the cDNA level (Goldstein et al. 2016; Kelemen et al. 2013; Shen et al. 2014; Trapnell et al. 2012; Vaquero-Garcia et al. 2016). Based on the isoform expression analyses, putative signal-specific functions of the isoforms were investigated. The expression of isoform MoYpd1p_T0 increases after treatment with KCl, whereas the expression of isoform MoYpd1p_T1 increases after treatment with sorbitol (Fig. 5). That indicates a role of MoYpd1p_T0 in salt stress response and MoYpd1p_T1 in sorbitol stress response. That is in line with the GFP studies, in which isoform MoYyp1p_T0 translocates into the nucleus after KCl stress, whereas MoYpd1p_T1 was found within the nucleus after sorbitol stress (Fig. 3). It was, therefore, hypothesized that the isoforms coexist and synergistically control stress responses and pathogenicity. Our hypothesis is supported by the transcriptomic analysis of field isolate KJ201, which detected infection-specific AS in about 500 genes (Jeon et al. 2022). However, *in vivo* studies on the biological function of isoforms of the pathogen are still needed. The role of MoB56 isoforms, for example, a regulatory subunit of protein phosphatase 2A, is not yet fully understood (Wang et al. 2023).

To conclude, this study provides a new model of the HOG signaling pathway *in M. oryzae* and underpins the major role of signaling pathways in fungal pathogenicity by adding the AS of signaling proteins as virulence determinants for the first time.

## 4. Material and Methods

### Fungal growth condition and sampling

The *M. oryzae* wildtype 70-15 and the *ΔMoypd1* mutant strain were routinely maintained, as described previously by Jacob et al. 2015. Regarding RNA isolation, conidia of 14-day-old M. oryzae cultures were harvested with H_2_O, filtered over two layers of Miracloth (Merck, Darmstadt, Germany), and used for the inoculation of 250 ml culture media (5 x 10^4^ conidia/ml) in a 500 ml glass flask. After three days of incubation at 26 °C and 120 rpm in a 12:12 light-dark cycle, an untreated sample was taken before the induction with stress (KCl [0.5 and 1 M], NaCl [0.75 M] sorbitol [0.5 and 1 M] and fludioxonil [10 μg/ml], respectively). Samples were then collected 25 and 180 min after stress exposure, frozen in liquid nitrogen and stored at – 80 °C.

### Generation of *YPD1*-fused GFP vectors

The process of isolating genomic DNA and transforming plasmids into NEB^®^ 10-β competent *Escherichia coli* strains was carried out as described previously (Bühring et al. 2021). Detailed information on the plasmid design and the primers required for the Gibson Assembly^®^, designed using the NEBuilder tool (https://nebuilder.neb.com/<u>#!/</u>), are provided in Figure 7 and

**Figure 7:**
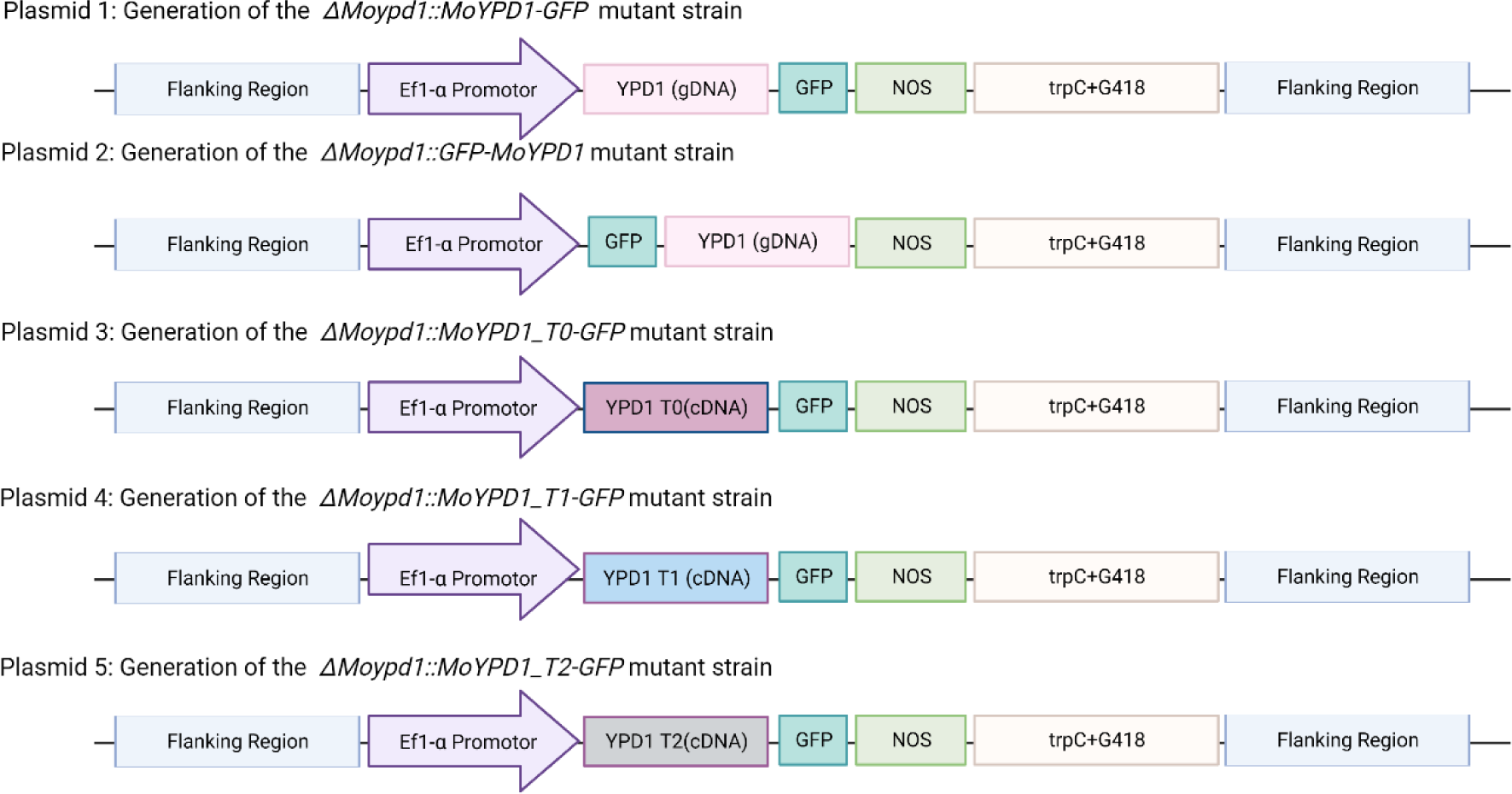
Overview plasmid design via Gibson Assembly^®^.

Table S1. MoYpd1p localization was investigated by mutants being transformed with plasmids in which either N-terminal or C-terminal GFP sequences have been fused to the genomic sequence of “total” *MoYPD1* (locus MGG_07173) or only to the CDS of the isoforms *MoYPD1*_T0 (MGG_07173T0) or *MoYPD1*_T1 (MGG_07173T1), as well as the recently discovered *MoYPD1*_T2 (Bühring et al. 2021). Total *MoYPD1*, 3’- and 5’-flanking sequences as well as the *Ef1*α promotor (of the gene MGG_03641, elongation factor 1 alpha) were amplified from gDNA whereas the single isoform-specific plasmids were generated by using cDNA as template. The remaining fragments were obtained from the *Agrobacterium*-compatible binary plasmid *pSJ + GFP*(G418) (Bohnert et al. 2019b), which served as the backbone for Gibson Assembly® after digestion with *Xho*I and *Bgl*II. The seven fragments of each plasmid 1–3 were assembled in one single reaction (Figure 7). Regarding the construction of plasmids 4 and 5, plasmid 3 was restricted with *XhoI* and *Bgl*II in order to introduce the respective isoform into the resulting backbone. The correct assembly was always confirmed by Sanger sequencing. Fungal transformation of the *M. oryzae* wildtype and the *ΔMoypd1* mutant strain was conducted using *Agrobacterium tumefaciens*-mediated transformation (Bohnert et al. 2019). The geneticin resistance cassette (G418) and the hygromycin resistance were used as resistance marker genes.

### Microscopy

Fluorescence microscopy of the GFP-fused *MoYPD1* strains was performed on the Revolution microscope (Echo) using Revolution software version 108.1. Conidia from 14-day-old fungal cultures grown on culture media agar plates were filtered through two layers of Miracloth (Merck) and incubated with Hoechst 33342 solution (20 mM, Thermo Fisher Scientific™) for 30 min at room temperature. The conidia were then mixed 1:1 with Biozym Low Melt Agarose (1.0 % w/v) and transferred to the 8-well chambers (ibidi ^®^). The NaCl [0.75 M], KCl [0.75 M and 1 M] or sorbitol [1 M] were added as stress-inducing agents.

### Plant infection assay

Conidia of the wildtype strain 70-15, *ΔMoypd1* and the respective *MoYpd1p* isoform-producing strains were harvested from 14-day-old cultures in order to assess virulence. Rice plants (CO39) were grown, as previously described, and treated by both drop and spray inoculation (Cao et al. 2022; Kramer, Thines, and Foster 2009).

### Vegetative growth and germination assay

Mutant strains were tested for stress tolerance on agar plates, as previously described by Jacob et al. 2014. In addition, germination was investigated according to Jacob, Schüffler and Thines 2016.

### RNA Isolation and cDNA synthesis

The lyophilized mycelium was mechanically disrupted using the TissueLyserII system (QIAGEN, Hilden, Germany) for 30 s at 30 Hz. Total RNA was extracted using the RNeasy^®^ Plant Mini Kit, including RNase-free DNase treatment, according to the manufacturer’s instructions. Three biological replicates of each condition were polled together. The RNA integrity numbers were evaluated by an Agilent 2100 bioanalyzer (Agilent Technologies, Santa Clara, CA, USA) employing the RNA 6000 Nano Kit (Agilent Technologies). Samples with RNA integrity number values above nine was selected for transcriptome analysis.

In order to detect new AS events by PCR and Sanger sequencing and plasmid construction, cDNA was synthesized with oligo(dT)20 primers with Invitrogen’s SuperScript^®^ III First-Strand Synthesis System for RT-PCR (Invitrogen, Germany), according to the manufacturer’s instructions. The cDNA used for ddPCR was transcribed using the iScriptTM Advanced cDNA Synthesis Kit for RT-qPCR (Biorad, Munich, Germany).

### PCR and ddPCR

Primer design (https://www.ncbi.nlm.nih.gov/tools/primer-blast/) was based on the results of Cufflinks, MAJIQ and SGSeq. The PCR was carried out in a total volume of 25 µL containing 1.25 µL of each primer [100 pmol/µL], 0.5 µL of dNTPs [10 mM], 2 µL of cDNA [100-600 ng/µl], 5 µL of 5 x Q5^®^ Reaction buffer, 0.5 µL of Q5^®^ DNA polymerase (NEB, Frankfurt, Germany), and 14.5 µL of RNase-free water to perform the reaction. Primer-specific annealing temperatures (Tm) were determined in advance using NEB Tm Calculator version 1.15.0, and extension times were calculated according to the manufacturer’s instructions. The PCR was carried out in a C1000 Touch Thermal Cycler (BioRad Laboratories, Hercules, CA, USA) under the following amplification parameters: initial denaturation for 30 s at 98 °C, 35 cycles consisting of denaturation for 10 s at 95 °C, annealing corresponding to primer-specific tm for 15 s, extension at 72 °C for the calculated extension time, and a final extension step for 5 min at 72 °C. The PCR product was separated by electrophoresis in 1 % agarose stained with ethidium bromide, and visualized using a QUANTUM-ST5-1100/26MX system (PEQLAB Biotechnologie GmbH, Erlangen, Germany). Amplicon gel extraction following the Monarch^®^ DNA Gel Extraction Kit (NEB, Frankfurt, Germany) instructions was used to isolate and purify the cDNA. Next, the GeneJET Plasmid Miniprep Kit (Thermo Fisher Scientific GmbH, Schwerte, Germany) was used to purify the plasmid DNA after cloning the PCR products using the CloneJET™ PCR Cloning Kit with DH10B Competent Cells (Thermo Fisher Scientific GmbH, Schwerte, Germany). Sanger sequencing was performed (Eurofins Genomics, Germany) to determine the sequence similarity between the amplified cDNA sequence and the *MoYPD1* sequence, followed by BLASTN analysis (https://blast.ncbi.nlm.nih.gov/Blast.cgi) and the prediction of ORFs (www.ncbi.nlm.nih.gov/orffinder).

We used the cDNA as a template for the ddPCR for the absolute quantification of a novel AS-Event. This technique is based on water emulsion droplet technology and fractionates the target template into 20,000 droplets. As a result, the PCR amplification of the template is carried out in each droplet, and counting the positive droplets allows for a precise, absolute determination of the target’s concentration. The ddPCR was performed using three independent replicates. A ddPCR reaction mixture of 11 μl QX200 EvaGreen Supermix (Bio-Rad Laboratories), 1.1 µl of each primer [2 pmol/µl], 1 µl cDNA [1000 ng/µl] and 7.8 µl water was used for the droplet formation. After droplet generation using the QX200 Droplet Generator (Bio-Rad Laboratories), 40 μl of sample was transferred to a 96-well plate (Bio-Rad Laboratories), which was heat-sealed at 180 °C for 5 s before amplification. The PCR was started by enzyme activation at 95 °C for 5 min, followed by 40 cycles of amplification at 95 °C for 30 s (denaturation) and 60 °C for 1 min (annealing/extension). After signal stabilization at 4 °C for 5 min and 90 °C for 5 min with a temperature ramp of 2.5 °C/s, the 96-well plate was incubated overnight at 12 °C. On the next day, an analysis was carried out using a QX2000 Droplet Reader (manufactured by Bio-Rad Laboratories). The reader utilizes Quanta Soft software, 1.2 standard edition (also manufactured by Bio-Rad Laboratories) in conjunction with a two-color detection system to analyze each droplet.

## Bioinformatic Analysis

### Protein Structure Prediction

The 3D protein structures of MoYpd1p_T0 (Uniprot ID: G4MTK9), MoYpd1p_T1 (Uniprot ID: L712312), MoSln1p (Uniprot ID: G4MV04) and MoHik3p (Uniprot ID: G4NKR9) were used to generate PPIs. Missing structures were predicted using AlphaFold version 2.3.2. The amino acid sequences required were obtained by Uniprot ID (Table S2). MoHik5p and MoHik6p could not be predicted due to the length of the amino acid sequences (1994 and 2580 amino acids, respectively).

### PPI Prediction

The PPIs between the MoYpd1p isoforms and HKs were modeled using predicted 3D structures by AlphaFold version 2.3.2 on the InterPred web server, available at the GRAMM Docking web server at https://gramm.compbio.ku.edu. Prediction occurred between each of the MoYpd1p_T0-T3 isoforms and the HK MoSln1p, Mohik1p-5p and Mohik7-9p isoforms.

### Sequencing

Illumina and PacBio Iso-Seq sequencing systems generated the RNA sequencing data in this study. Library preparation and Illumina 100 bp paired-end sequencing of total RNA were performed by Eurofins Genomic Europe Sequencing GmbH (Germany), followed by a raw data quality assessment with QualiMap version 2.2. (https://github.com/EagleGenomics-cookbooks/QualiMap). Depending on the analysis tool, TopHAT (Galaxy version 2.1.1) and HISAT2 (Galaxy version 2.2.1+ galaxy1) aligned the raw FASTQ data derived by short-read sequencing along the 70-15 reference strain genome (assembly version MG8, http://fungi.ensembl.org/Magnaporthe_oryzae/Info/Index). Furthermore, full-length transcript analysis was performed using PacBio’s Iso-Seq method at the Berlin Institute of Health at Charité (Germany). After Iso-Seq library preparation and sequencing on the PacBio Sequel II platform (one SMRT^®^ Cell 8M), raw reads were classified into circular consensus sequences and non-circular consensus sequences subreads and processed as described in IsoSeq3 pipeline version 3.8.1 (https://isoseq.how/). To this end, the Pigeon workflow was used for mapping, merging and classifying transcripts according to the IsoSeq3 cluster of the bulk workflow.

### Alternative splicing analysis of *MoYPD1*

Several approaches, including isoform-based (Cufflinks/Cuffdiff) and splicing event-based (rMATS, MAJIQ, and SGSeq) analysis, were conducted for the prediction of *MoYPD1* isoforms. The expression values were determined using StringTie and Ballgown and DESeq2.

### Cufflinks

Cufflinks, as an isoform-based method, involves several programs, including Cufflinks itself, Cuffmerge and Cuffdiff (Fig. 8). Cufflinks version 2.2.1 was used, following the instructions available at http://cole-trapnell-lab.github.io/cufflinks/manual/. FASTQ files of two experimental conditions (with and without stress induction) were mapped to MG8 reference annotation using TopHat. Based on the resulting SAM or BAM file, Cufflinks assembled the alignment files independently of the reference annotation to possible transcripts and generates transcriptome assemblies for each condition (GTF files). The assemblies of each condition and the MG8 reference annotation were used for merging by Cuffmerge, resulting in a final transcriptome assembly (GTF file). Following this, TOPHAT-based BAM files and the final transcriptome assembly (GTF file) were used for isoform-based analysis with Cuffdiff. Finally, the R package cummerbund version 2.38.0 was used for visualization; it reads the output files from Cuffidff into an S4 object, making the data easily accessible for graphical representation.

**Figure 8:**
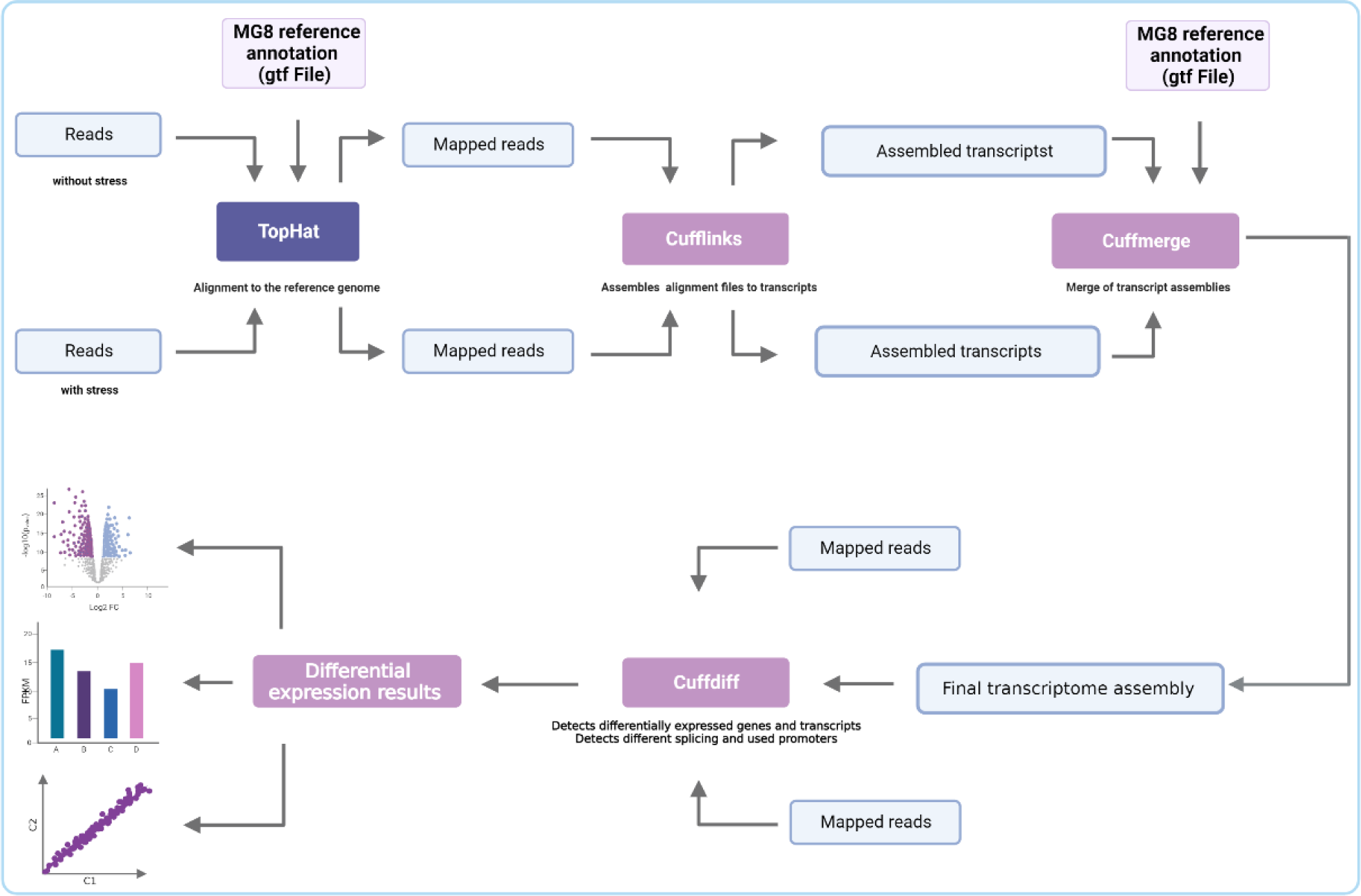
Workflow of RNA Seq analysis via Cufflinks.

### rMATS, MAJIQ and SGSeq

RNA-Seq data were also analyzed with HISAT2-produced alignments and count-based methods. The MG8 annotations were processed to generate AS events using rMATS turbo version 4.2.1, as described at https://github.com/Xinglab/rmats-turbo/blob/v4.1.2/README.md. Data were filtered with a cut-off p-Value of 0.05. In contrast to Cufflinks, rMATS provides explicit AS events rather than whole transcripts. The identification of known and *de novo* splice connections were performed using SG-Seq and MAJIQ. Local splicing variations were analyzed with MAJIQ version 2.4, conducted as described at https://biociphers.bitbucket.io/majiq-docs-academic/. Local splicing variations can be visualized as splits in splice graphs where several edges are connected to or derived from a single exon, called the reference exon (Vaquero-Garcia et al. 2016). Based on the aligned reads, the MAJIQ Builder constructed a splice graph for *MoYPD1,* then local splicing variations for each condition are quantified and compared with the MAJIQ Quantifier, calculating the percent inclusion index (Ψ) and its change (ΔΨ). Finally, the Viola visualization package displayed the results of the MAJIQ Builder and Quantifier as a splice diagram with alternative splice variants and violin plots of *MoYPD1* depicting the Ψ and ΔΨ estimates. The AS events were also identified with SGseq version 1.30.0, an R/Bioconductor package (https://bioconductor.org/packages/release/bioc/vignettes/SGSeq/inst/doc/SGSeq.html). Mapped reads are used to predict splice junctions and exons in a genome-wide splice graph. Based on reads extending across the start or end of each splice variant, recursive splice events are identified on the graph and quantified locally (Goldstein et al. 2016).

### Sashimi plot

The visualization of splice junctions from HISAT2-aligned RNA-seq data along the *MoYPD1* gene of reference strain 70-15 was performed as described at https://github.com/guigolab/ggsashimi. Arcs representing introns include the number of reads spanning an exon in Sashimi plots. Only splice junctions with a minimum read coverage of 15 for *MoYPD1* are shown.

### StringTie and Ballgown

Heat plots for transcript expression were created using StringTie version 2.2.1 and Ballgown version 2.28.0. StringTie, firstly, builds AS graphs and then performs a heuristic algorithm to determine the heaviest path, which represents a transcript. Afterwards, a flow network design is used to determine the coverage of each transcript (Pertea et al. 2015). Here, the Ballgown compatible output was generated providing a HISAT2-generated alignment and the MG8 annotation (GTF file) for StringTie, and was used subsequently to visualized the transcripts and their expression level under different conditions, as described at https://github.com/alyssafrazee/ballgown.

### DeSEq2

Salmon version 1.9.0 (https://salmon.readthedocs.io/en/latest/) was used for the initial quantification of transcripts. The differential expression analysis was conducted using version 1.36.0 of DESeq2 (Differential Expression Analysis for Sequencing Data 2), according to http://www.bioconductor.org/packages/release/bioc/vignettes/DESeq2/inst/doc/DESeq2.html.

## Acknowledgments

We are grateful for support by Prof. Dr. Elmar Jaenicke, Institute of Molecular Physiology (IMP), Johannes Gutenberg-University, Mainz, Germany in predicting protein structures. In addition, the technical support provided by Jessica Behnke is greatly appreciated.

